# Speed Kills: Accelerated Pharmacokinetics Reduce Persisters in *Escherichia coli*

**DOI:** 10.1101/2025.04.25.650648

**Authors:** Khairy M Ali, Ronan A. Murphy, Caroline Zanchi, Sara Zunk Parras, Alexandro Rodriguez-Rojas, Jens Rolff

## Abstract

Persister cells are bacterial cells that form small subpopulations and are refractory to antimicrobials. Importantly, those cells can cause the relapse of an infection and also provide a starting point for antimicrobial resistance evolution. By now, several mechanisms of persister formation have been studied. For example, it has been shown that a short exposure to sublethal concentrations of antimicrobials can induce persister formation. Given this, we hypothesised that the pharmacokinetics, the temporal changes of drug concentrations under treatment, could impact persister formation. We predicted that the longer bacteria spend under sublethal drug concentrations, the more persisters would form. Using a set-up with small chemostats and the antimicrobial peptide Pexiganan, we indeed found that faster pharmacokinetics resulted in lower persister numbers compared to slower pharmacokinetics. This finding provides a proof of principle that pharmacokinetics, which can be influenced by treatment procedures, has consequences for persister formation. Our results suggest that faster pharmacokinetics could be useful to minimize persister numbers and hence to lower the risks of relapse or resistance evolution.

## Introduction

Persister cells, a subpopulation of bacteria that are refractory to antimicrobials without change in their antimicrobial susceptibility, are thought to be a major factor in the persistence of chronic infections. This is because antimicrobials typically eliminate most bacterial cells but persisters survive (Fisher et al., 2017a; Stojowska-Swędrzyńska et al., 2023). Persisters were first described by Hobby et al. (1942) who demonstrated that penicillin was ineffective against non-growing *Staphylococcus aureus* cells, indicating a connection between persister cells and dormant states. Since then, accumulating experimental evidence supports that dormancy in persister cells is characterized by a lack of growth, as well as the absence of transcription, translation, and proton-motive force, rendering most antimicrobials ineffective (Bigger, 1944; Fisher et al., 2017; Hobby et al., 1942; Kwan et al., 2013). Persister cells can cause a relapse of infection when antimicrobial concentrations decrease and can lead to the evolution of antimicrobial resistance (Balaban et al., 2004, 2019; Fisher et al., 2017). Bacterial populations that contain persisters are described by a biphasic killing curve that shows a small subpopulation which dies at much slower rate than the overall population (Balaban et al., 2019).

While Persisters emerge spontaneously within bacterial population, their formation can be induced. Known triggers for persister formation include, heat stress, pH stress, oxidative stress and antimicrobials (among others) (Niu et al., 2024). By now several mechanisms and genes have been identified that lead to persister formation. We found in a previous study that priming (short sub-lethal exposures) of *Escherichia coli* by two antimicrobial peptides (AMPs) - melittin and pexiganan - increases persister formation leading to subsequent greater survival to lethal doses of the same AMPs (Rodríguez-Rojas et al., 2021).

AMPs are found across all classes of life and serve as a fundamental component of innate defence mechanisms against microbial pathogens. They are promising as new antimicrobials and currently constitute 15 % of all antimicrobials on the World Health Organisation’s (WHO) list of antimicrobial drugs under preclinical trials (2024). In innate immunity, AMP production starts at sub-lethal concentrations and can take several hours to reach lethal levels upon infection (Makarova et al., 2016). The same is also true for clinical antimicrobial treatments where this increase of drug concentration (its pharmacokinetics) takes time and goes through a concentration gradient.

In the light of these facts, and our finding that a brief sublethal exposure to AMPs induces persister formation, our study aims at exploring whether the longer a population of bacterial cells is exposed to sublethal concentrations of antimicrobials, here AMPs, the more persisters cells will form. This approach highlights the effect of pharmacokinetics, where drug concentrations increase gradually which resembles clinical applications (e.g., during antimicrobial infusion or localized antimicrobial delivery). This could improve our understanding of the recurrent bacterial infections after treatment courses in a clinical setting.

We used a chemostat system to explore how prior exposure to sub-lethal doses of the AMP pexiganan promotes bacterial survival through persistence and how different pharmacokinetics affect the induction of persistence. The chemostat system allowed us to dynamically control the concentration of the stressor, a critical advancement over previous studies that relied on static concentrations (Rodríguez-Rojas et al., 2021). By increasing the pexiganan concentration at different rates, we were able to investigate how varying exposure rates influence persister formation. This new approach provides deeper insights into how bacteria adapt over time, making it particularly relevant for understanding the development of antimicrobial persistence.

## Methods

### Bacterial strain

*E. coli* MG1655 was used for all experiments. The strain was stored at -70°C in a glycerol suspension and revived onto lysogeny broth agar (LB agar) (Carl Roth, Germany). Overnight liquid cultures were performed in Mueller Hinton Broth (MHB) (Carl Roth, Germany) in 50 mL falcon tubes.

### AMP Priming

Priming with pexiganan was performed according to a protocol of Rodríguez-Rojas et al., (2021). Briefly, overnight bacterial cultures were diluted 1:100 into fresh MHB in 50 mL tubes and incubated until cultures reached mid-log phase (OD_600_ 0.5, ∼ 2 × 10^8^ CFU/mL); ∼2 hr. Bacterial cultures were exposed to 0.4 µg/mL pexiganan (MIC/10) for 30 min at 37°C with shaking. Non-primed cultures were incubated in parallel. Following incubation, bacteria were washed by centrifugation at 4000 x g for 10 min, supernatant removed and resuspended in an equal volume of fresh MHB. Subsequently, cultures were allowed to recover for 60 min at 37°C with shaking.

### Chemostat system and dynamic application of AMPs

After recovery, 3 mL of primed and non-primed cultures were loaded into sterile 5 mL chemostat tubes, containing a magnetic stirrer. The chemostats were placed on a magnetic stirrer (Variomag Multipoint HP 6) and connected to a syringe pump (Braintree Scientific Programmable Syringe Pump, Braintree Scientific, Inc. USA). Syringes containing pexiganan in MHB (details below) were connected to the chemostats by peristaltic tubes. The chemostat system was placed inside an incubator at 37°C (Figure 1A).

**Figure 1.**
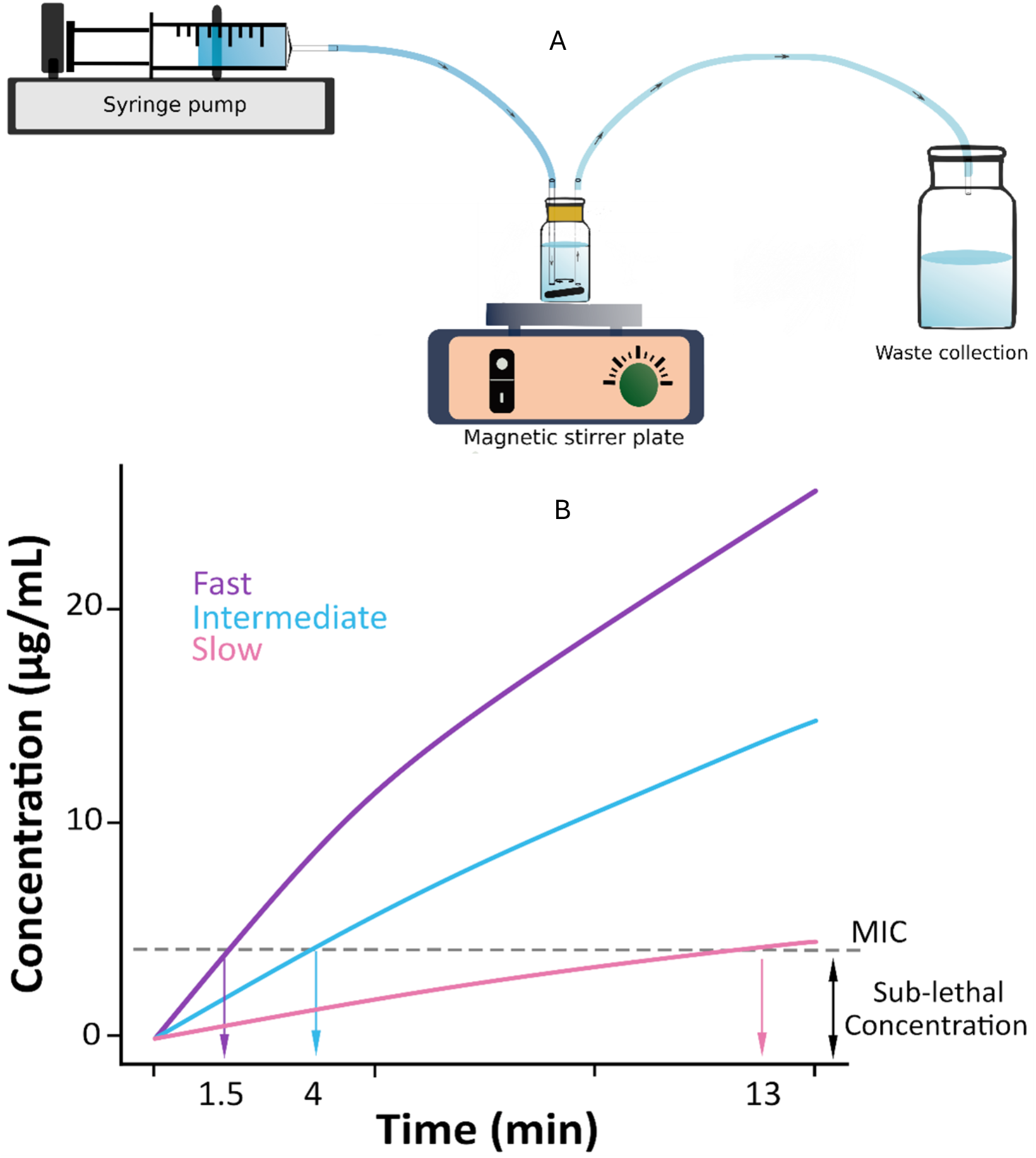
(A) Chemostat system inside an incubator at 37oc, a syringe pump delivers pexiganan into a chemostat tube positioned on a magnetic stirrer plate to ensure homogeneous mixing. The bacterial suspension/pexiganan is simultaneously withdrawn from the vessel into a waste collection container. Arrows indicate the direction of flow through the Chemostat system. (B) Pharmacokinetic of AMP concentration increases in the chemostat systems. PEX was applied in three distinct dynamics: **Slow** (concentration 40 µg/mL, rate 25 µL/min), **Intermediate** (concentration 80 µg/mL, rate 45 µL/min), and **Fast** (concentration 80 µg/mL, rate 100 µL/min). As a result, the PEX MIC for E. coli MG1655 was reached at 13 min for **Slow**, 4 min for **Intermediate**, and 1.5 min for **Fast**.

Cultures were challenged with pexiganan supplied in three distinct application regimes (Figure 1B. Figure S1): for the *Slow* application, 40 µg/mL was applied at a rate of 25 µL/min (MIC reached after 13 min), for the *Intermediate* 80 µg/mL at 45 µL/min (MIC reached after 4 min), and for the *Fast* 80 µg/mL at 100 µL/min (MIC reached after 1.5 min). Ten µL of samples were collected using a 1 mL syringe, serially diluted in phosphate buffered saline (PBS) and spot-plated onto LB agar, to determine CFU/mL at time points 2.5, 5, 7, 15, 30, 60, 90, 120 min. Pumping was stopped at 120 min and a final sample collected 30 min later. In parallel to each run, a control was run and sampled consisting of a mid-log bacterial culture connected to a syringe containing MHB-only (no prime, no challenge). The bacterial load in these tubes did not differ between the 3 application regimes (Figure S2) or across the duration of the experiments (Chisq = 1.4418, *p* = 0.4863), indicating that any differences in CFU/mL observed subsequently were not due to the chemostat conditions or sampling procedure.

### Persister survival assay

To determine the fraction of persisters present in a given culture from the *Slow, Intermediate*, and *Fast* application regimes (at time points with countable colonies), we followed the protocol by Shan et al. (2017): a 4 hr exposure to ciprofloxacin (CIP) at 2 μg/mL was performed at 37°C. After exposure, bacteria were washed twice with 0.9% NaCl, serially diluted and plated on MHB agar. The bacterial density immediately before the addition of CIP was also determined (i.e. timepoint = 0). The fraction of persisters was calculated as [bacterial number after CIP treatment / bacterial number before CIP treatment] (Rodríguez-Rojas et al., 2021).

### Statistics

All statistical analyses were performed using the R version 4.4.1 (R Core Team, 2024). We first analysed whether priming increased the fraction of persisters in bacterial cultures. To do so, the ratio of persisters/total population size was analysed as a response variable with a beta-regression (‘betareg’ package, (Kosmidis & Zeileis, 2024)). The priming treatment (primed – unprimed) was the explanatory variable.

All figures were generated using (‘ggplot2’ package,(Hadley Wickham, 2016)). To samples with zero colony-forming units (CFUs), a value of 1 was added to all final CFU counts prior to visualization.

The evolution of the bacterial population size over time between the primed and unprimed treatments under the 3 different dynamics was analysed in 2 steps. Because there were samples from which we could not retrieve viable CFU/mL, we first analysed the presence/absence of CFU/mL in samples with a Generalized Linear Mixed Model (GLMM) fitted for a binomial distribution using ‘glmmTMB’ function of the ‘glmmTMB’ package, according to time/concentration, treatment, and the interaction between them as explanatory variables (Brooks et al., 2017). The starting bacterial culture was included as a random factor.

We then focused on samples from which we could retrieve viable CFU/mL. In these, the number of CFU/mL was analysed with a GLMM fitted for a negative binomial distribution, according to time/concentration and treatment as well as the interaction between them, plus the starting bacterial culture as a random factor (Brooks et al., 2017).

Last, we analysed whether the proportion of persisters differed according to the dynamics considered. To do so, the ratio of persisters: total population size was analysed as a response variable at 15, 60 and 120 minutes of each pexiganan application regime (where possible). We then analysed the proportion of persisters separately for each time point with a beta regression (‘betareg’ package) according to the application regime and the priming treatment as explanatory variables, as well as the interaction between them. This means that at 15 minutes, the data relative to the Slow, Intermediate and Fast are being analysed. At 60 minutes, the data of Slow and intermediate are analysed, whereas at 120 minutes, only the data relative to the Slow dynamics are analysed, according to the priming treatment.

The effect of the explanatory variables which were significantly affecting the response variables are represented in the Supplementary Material as plots of the estimates with 95 % Confidence Intervals (95CI), which allows the reader to perform pairwise comparisons between treatment levels by eye. The difference between 2 treatment levels is deemed significant when 95CI do not overlap more than half of their length (Cumming, 2009; Cumming & Finch, 2005).

## Results

### Generation of persisters by priming

Persisters were detected by treating pexiganan primed/non-primed cultures for 4 hr with CIP. We analysed whether the proportion of persisters of MG1655 changed when the culture was primed or not (Figure S3) (Rodríguez-Rojas et al., 2021). As expected, primed cell cultures produced a higher persister proportion than non-primed cell cultures (Chisq = 4.3132, *p* = 0.03782).

### *Slow* AMP application

We found neither an interaction between time/concentration and priming (Chisq = 0.0300, *p* = 0.8) on the proportion of surviving populations, nor an effect of priming alone (Chisq =0.0052, *p* = 0.9). Only time/concentration significantly affected the proportion of surviving populations in the *Slow* application regime (Chisq = 6.1498, *p* < 0.005). The proportion of surviving populations decreased with time/concentration (Figure 2)

**Figure 2.**
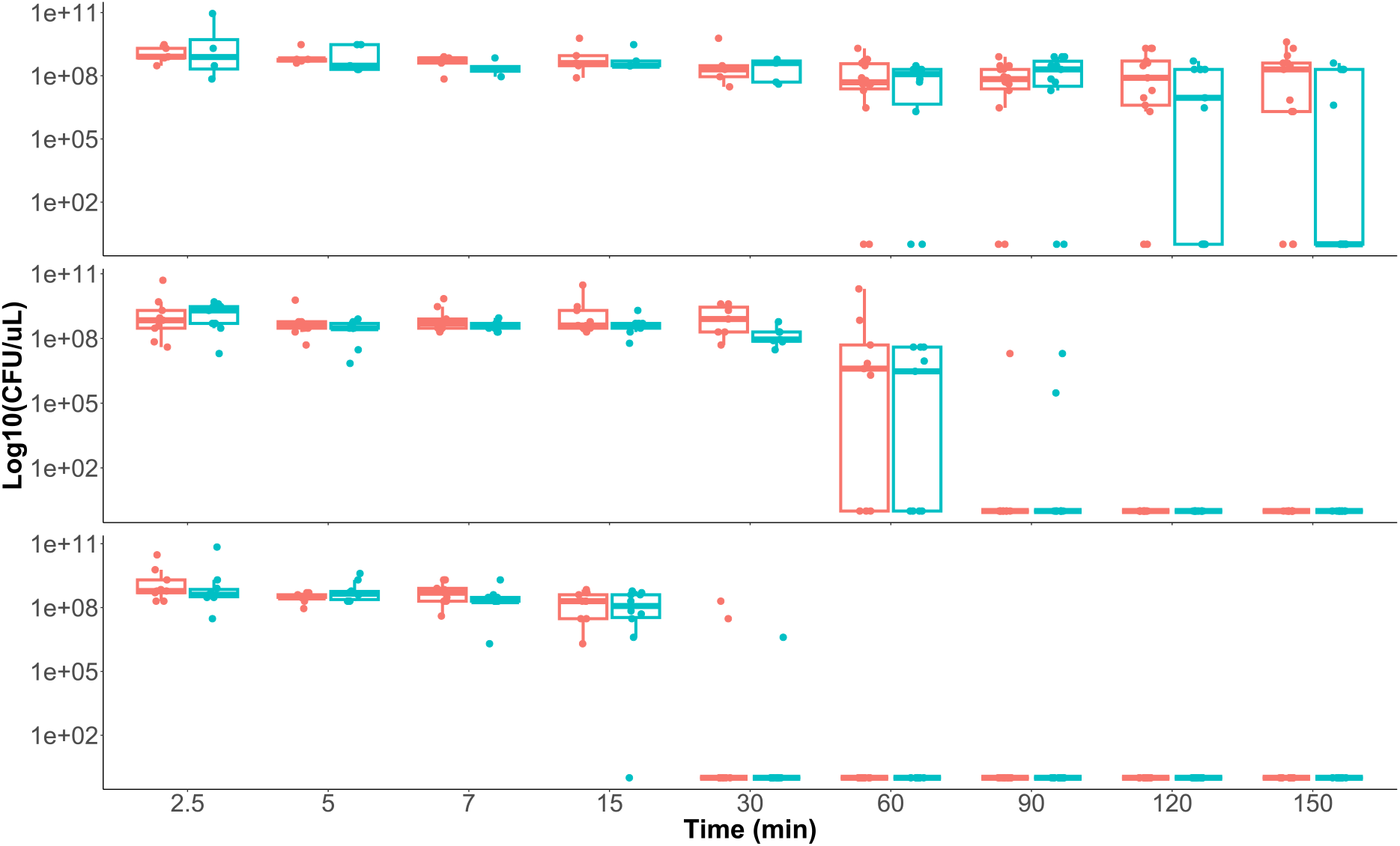
Survival of primed/non-primed populations under dynamic application of pexiganan. **Slow** (top), **Intermediate** (middle) and **Fast** (bottom)dynamics are shown. The red boxes and dots represent the results of the non-primed bacterial cultures, whereas the blue boxes represent the result of primed bacterial cultures. For raw CFU values check Figure S4.

In the populations that survived, there was no significant effect of treatment, neither in interaction with time/concentration (Chisq = 3.8632, *p* = 0.5693) nor as a simple effect (Chisq = 0.0722, *p* = 0.7881) (Figure S5). The number of CFU/mL in the samples significantly decreased over time/concentration (Chisq = 83.2032, *p* <0.005).

### 2.2. *Intermediate* AMP application regime

Similarly to the *Slow* application regime, the proportion of surviving populations was not affected by the interaction between time/concentration and treatment (Chisq = 1.7512, *p* = 0.1857) or the priming treatment alone (Chisq = 2.6551, *p* = 0.1032). The proportion of surviving populations did significantly decrease with time/concentration (Chisq = 1.3864, *p* = 0.2390): most of the populations present in primed as well as non-primed samples had collapsed to zero CFU/mL at 90 min (37 µg/mL) (Figure 2).

The result is similar in the populations that survived, where there was no significant effect of treatment, neither in interaction with time/concentration (Chisq = 1.7637, *p* = 0.1842) nor as a simple effect (Chisq = 0.9679, *p* = 0.3252). The number of CFU/mL in the samples significantly decreased over time/concentration (Chisq = 40.3886, *p* <0.005).

### 2.3. *Fast* AMP application regime

The populations in 7 out of 9 primed and 9 out of 10 non-primed samples had collapsed to zero at 30 minutes (34 µg/mL). This rendered the analysis of the proportion of surviving populations over time irrelevant. Hence, we analysed the number of CFU/mL in populations that survived up to 15 min (Figure 2. Figure S5).

There was again only a significant effect of time/concentration (Chisq = 13.1786, *p* < 0.005) on the number of CFU/mL retrieved from the samples, but neither a significant interaction between time/concentration and priming (Chisq = 2.2536, *p* = 0.521477), nor an effect of priming (Chisq = 0.2163, *p* = 0.641894) (Figure 2).

When comparing the 3 application regimes, we noticed that in the *Slow*, while the populations present in some samples do collapse starting from 60 minutes, the subsequent time points comprise a mixture of populations containing CFU/mL and collapsed populations (2 out of 14 in primed collapsed and 2 out of 11 in non-primed). In the *Intermediate* regime, the drop in the proportion of samples containing viable CFU/mL is more dramatic, since the populations present in 8 out of 9 primed and 7 out of 9 non-primed samples had collapsed to zero CFU/mL at 90 min (37 µg/mL).

## 3. Presence of persisters in different application regimes

We tested how the proportion of persisters vary with time spent under different application regimes of pexiganan to MG1655 (Figure 3).

**Figure 3.**
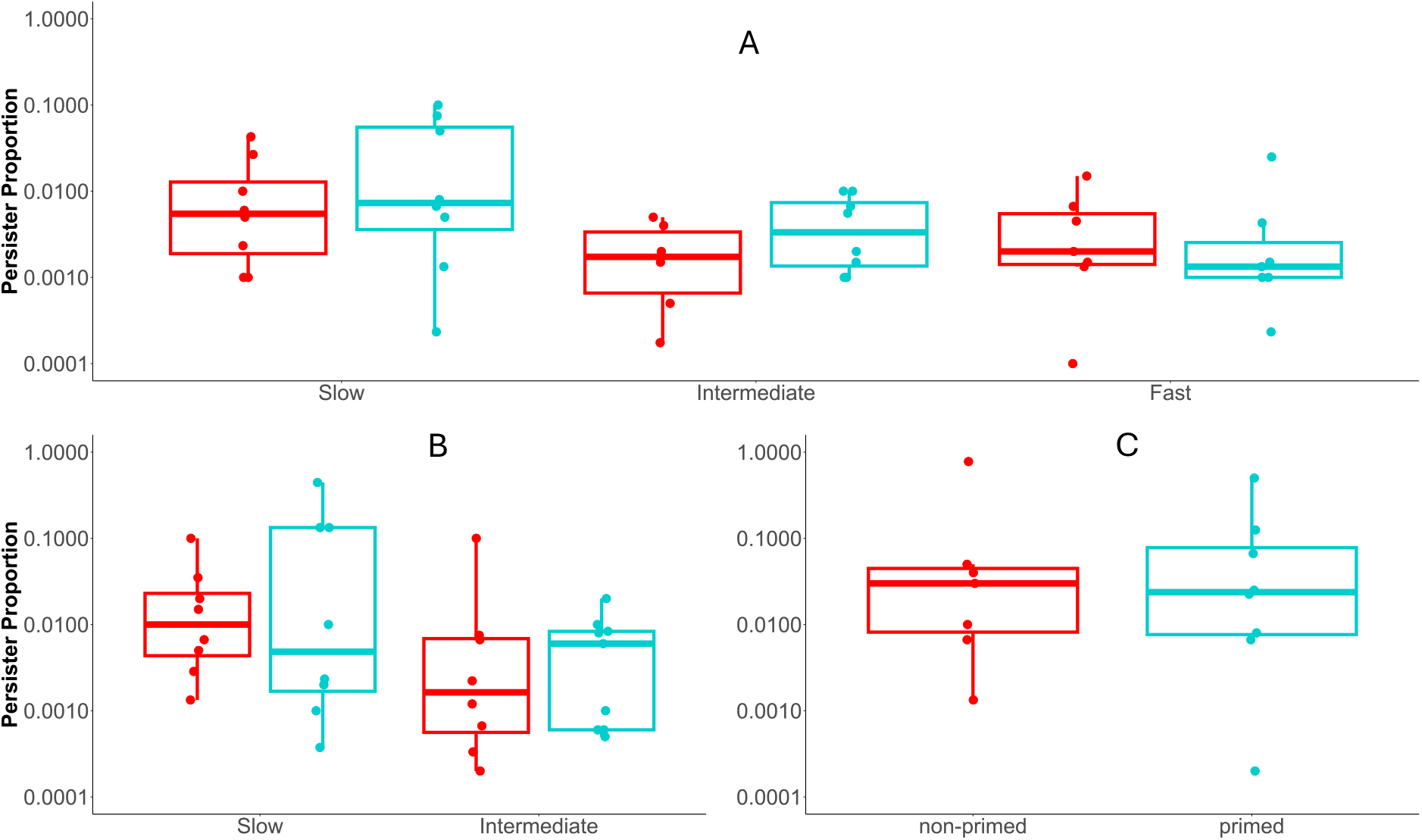
Persister cell proportions and estimated CFU/mL Under different pexiganan application regimes. The red boxes and dots represent the results of the non-primed bacterial cultures, whereas the blue boxes and dots represent the results of the primed cultures. (A) 15 min: persister proportion in **Slow, Intermediate** and **Fast** application regimes, the point of collapse for **Fast** regime. (B) 60 min: persister proportion in **Slow, Intermediate** application regime, point of collapse of **Intermediate** regime. (C) 120 min: only **Slow** regime had viable persisters.

We sampled cell cultures at 15, 60 and 120 minutes of each pexiganan application (where possible) and assessed the proportion of persisters with CIP. We analysed the effect of priming and the application speed on the proportion of persisters separately for the 3 time points sampled. Since at 15 minutes, the *Slow, Intermediate* and *Fast* application regimes all contained viable cells, the analysis of this time point compares the proportion of persisters from all 3 application regimes. At 60 minutes however, we could only retrieve viable CFU/mL in the *Slow* and *intermediate* application regimes, limiting the comparison of the proportion of persisters found in these 2 regimes. The 120 min time point analysis was performed on cells from the Slow application regime only.

At 15 min, (*Slow, Intermediate* and *Fast* application regimes), we found there was a significant effect of the application regime (Chisq = 7.7443, *p* <0.005). The *Slow* AMP application regime yielded significantly more persisters than the *Fast* one (Figure S6). However, the proportion of persisters found in the *Intermediate* application regime was not significantly different from the proportion of persisters found in the *Slow* and *Fast* regimes. There was again no effect of priming on the proportion of persisters, neither in interaction with application regime (Chisq = 0.5904, *p* = 0.7444), nor as a single effect (Chisq = 1.4812, *p* = 0.22359).

At 60 minutes, however (*Slow* and *Intermediate* application regimes), the proportion of persisters was neither explained by the interaction between priming and application regimes (Chisq = 0.1406, *p* = 0.7077), nor by the simple effects of priming treatment (Chisq = 0.3258, *p* = 0.5682) or application regime (Chisq = 2.945, *p* = 0.08614).

The priming treatment also had no significant effect on the proportion of persisters at 120 minutes (*Slow* application regime only, Chisq = 0.0294, *p* = 0.8639167)

## Discussion

The results shown here demonstrate the relationship between bacterial survival, persister formation, and AMP pharmacokinetics. To our knowledge, this study is the first to use chemostats to test the effect of AMPs on persistence in bacteria, an effort to model more closely the manner in which bacteria are exposed to antimicrobials in natural settings. Testing priming with a dynamic approach drug application, showed a contrast with previous observations where priming provided a survival advantage against a static pexiganan exposure at a lethal concentration. In this case, while priming with pexiganan did increase the proportion of persisters in bacterial populations, this effect did not translate in a higher bacterial survival in the 3 different application regimes of pexiganan.

Despite finding differences between persister formation in different application regimes, time points, assays and analyses, no differences in the proportion of persisters were ever seen between primed and non-primed bacterial populations. In static experiments, the AMP challenge by pexiganan and melittin (10 X MIC) imposed on the population was sudden and fixed, giving the previously exposed, primed bacteria an advantage over non-primed cells (Rodríguez-Rojas et al., 2021). By contrast, with the gradual increase of AMP concentration within the chemostat system, non-primed populations are not suddenly challenged with a lethal concentration but rather also pre-exposed to sub-MIC concentrations (primed) as the AMP increases gradually. This most likely negates the priming advantage by providing an opportunity for non-primed populations to catch up to primed populations in their ability to form persisters, i.e. priming is unavoidable for non-primed during dynamics application of pexiganan.

We saw that a faster increase in the concentration of pexiganan increased the rate of bacterial death: from 30 min onwards *Fast* pexiganan application rendered the majority of the populations non-viable, whereas all replicates in the *Intermediate* and *Slow* application regimes still contained viable cells at this time point. From 90 min onwards in the *Intermediate* application regime the majority of replicates were not viable whereas in the *Slow* application regime the majority of cells were viable throughout the duration of the experiment (150 min). We could retrieve more persisters in the *Slow* than in *Intermediate* and *Fast* application regimes of pexiganan at 15 minutes into the experiment, time point by which the samples of these 3 application regimes are all past MIC.

Pharmacokinetic/pharmacodynamic models (PKPD) explain resistance evolution and treatment failure by showing how antimicrobial dynamics shape the bacteria’s survival (Yu et al., 2018). Consistent with the modelling approach reviewed by (Witzany et al., 2023), which has demonstrated that the rate of drug application significantly influences bacterial population dynamics, we observed a decline in the overall population sizes from the start of treatment under all three dynamics. The extent of this decline is directly influenced by the rate of the drug application. Under Fast pharmacokinetics, there is a drastic increase in concentration, translated into a short survival period for the overall populations and resulting in fewer persisters. In contrast, Slow pharmacokinetics allowed for prolonged bacterial survival and higher persister numbers due to extended sub-MIC exposure. The different pharmacodynamics not only led to different numbers of persisters, with the highest number forming under the Slow regime, but also to a shorter time window when persisters can actually form.

Our finding of a relationship between rate of AMP application and bacterial survival rates during pharmacokinetic treatments could also be (partially) explained by the inoculum effect. The inoculum effect is the phenomenon where the higher the bacterial population density the higher the drug concentration required to kill that population, and is particularly pronounced for AMPs (Baeder & Regoes, 2019; Loffredo et al., 2021; Seeger et al., 2021). Our *Slow* pexiganan increase extends the time spent sub-MIC, allowing high numbers of cells to survive and proliferate, thereby maintaining a high population density. This elevated density amplifies the inoculum effect, which raises the effective MIC required for eradication. As a result, even when pexiganan levels surpass MIC, the surviving population remains dense enough for long enough to resist killing by the AMP, while the prolonged sub-MIC stress simultaneously promotes transitions into the persister state; a vicious cycle of greater cells numbers negating concentration increases and fueling persister formation. In contrast, *Fast*/*Intermediate* regimes rapidly exceed MIC, limiting the duration of sub-MIC exposure and hence both population growth and persister formation initiating virtuous cycles of reduced population densities, higher AMP concentrations and a reduced reservoir for the switch to bacterial persistence.

Our results here are a proof of principle that exposure time to sub-lethal concentrations of antimicrobials caused by different pharmacokinetics has consequences for persister formation. The slower the increase in drug concentration, the more persisters that are formed. If this finding were to be repeated in other drug/bacteria combinations, this effect has potentially important implications for treatments. Given that persisters can cause the relapse of infection, reducing the proportion of persisters should be a goal of treatment (Balaban et al., 2019; Fauvart et al., 2011). The results presented here indicate that faster pharmacokinetics are desirable as they are better at reducing bacterial population densities and at preventing persister formation than slower pharmacokinetics and pharmacokinetics should be a strong consideration in antimicrobial administration in infection treatment.

