## Supplement figures for "Speed Kills: Accelerated Pharmacokinetics Reduce Persisters in *Escherichia coli*"

### Supporting information

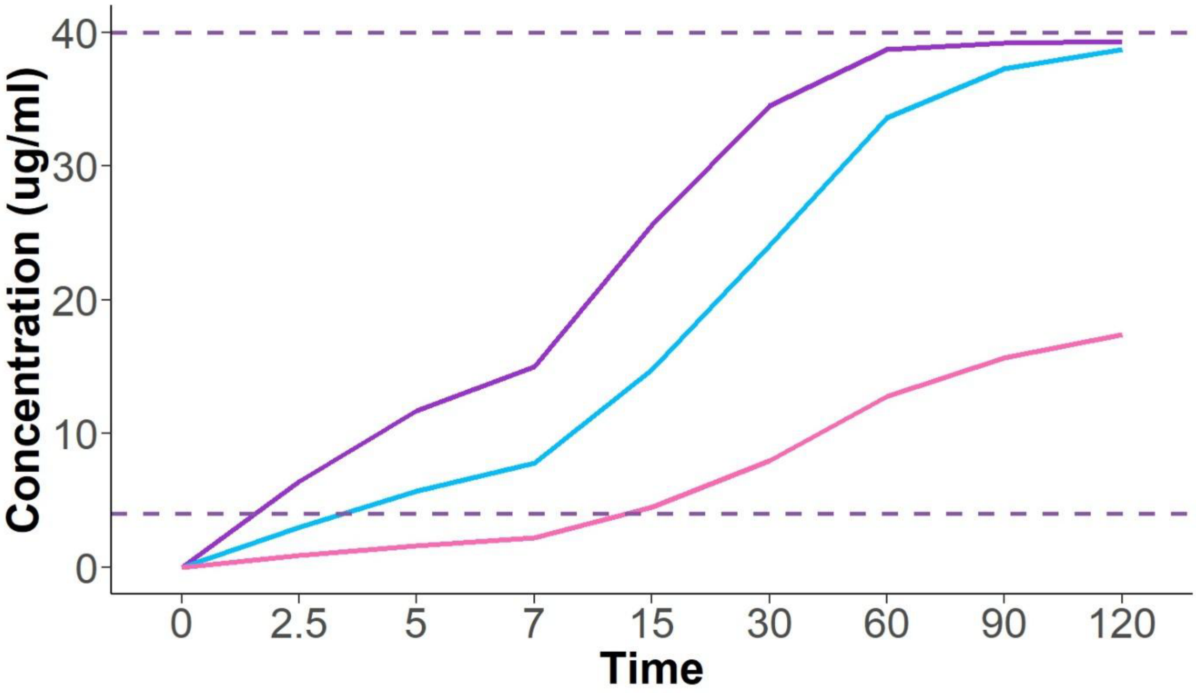


*Figure S1A Pharmacokinetic rates of pexiganan in the chemostat system. Purple line represents the **Fast** dynamic, blue line is the **Intermediate** dynamic, and pink is the **Slow** dynamic. Dashed line in the bottom is the MIC and on top is 10x MIC.*

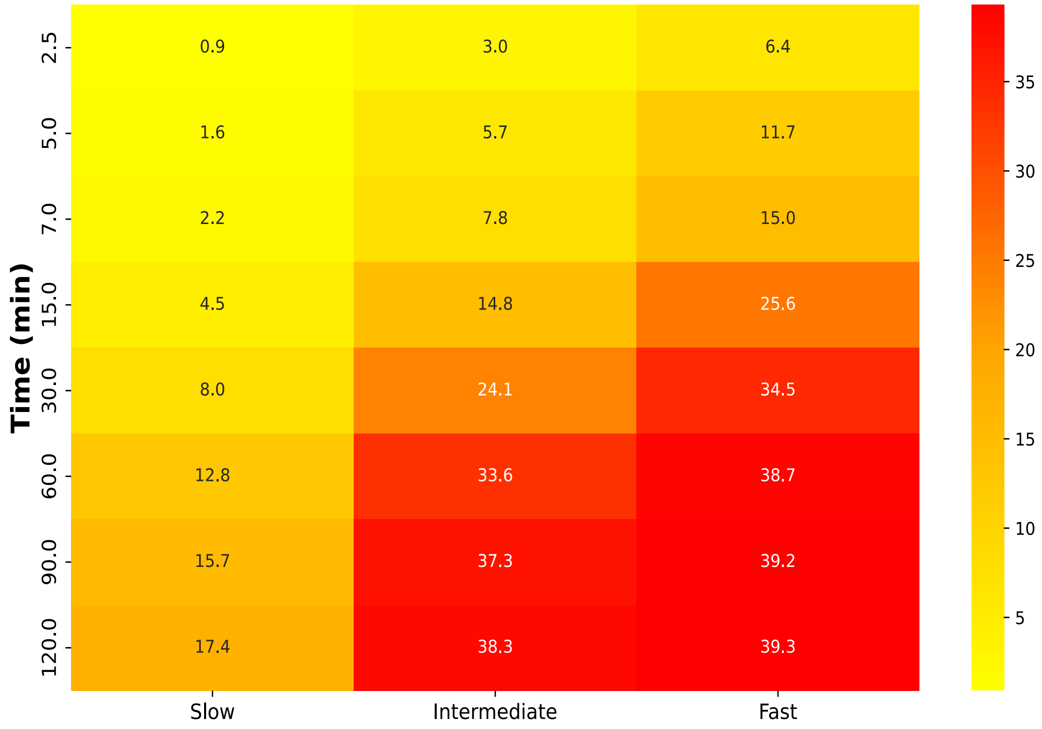


*Figure S1B Heat map of pharmacokinetic increase rates of pexiganan in the chemostat system.*

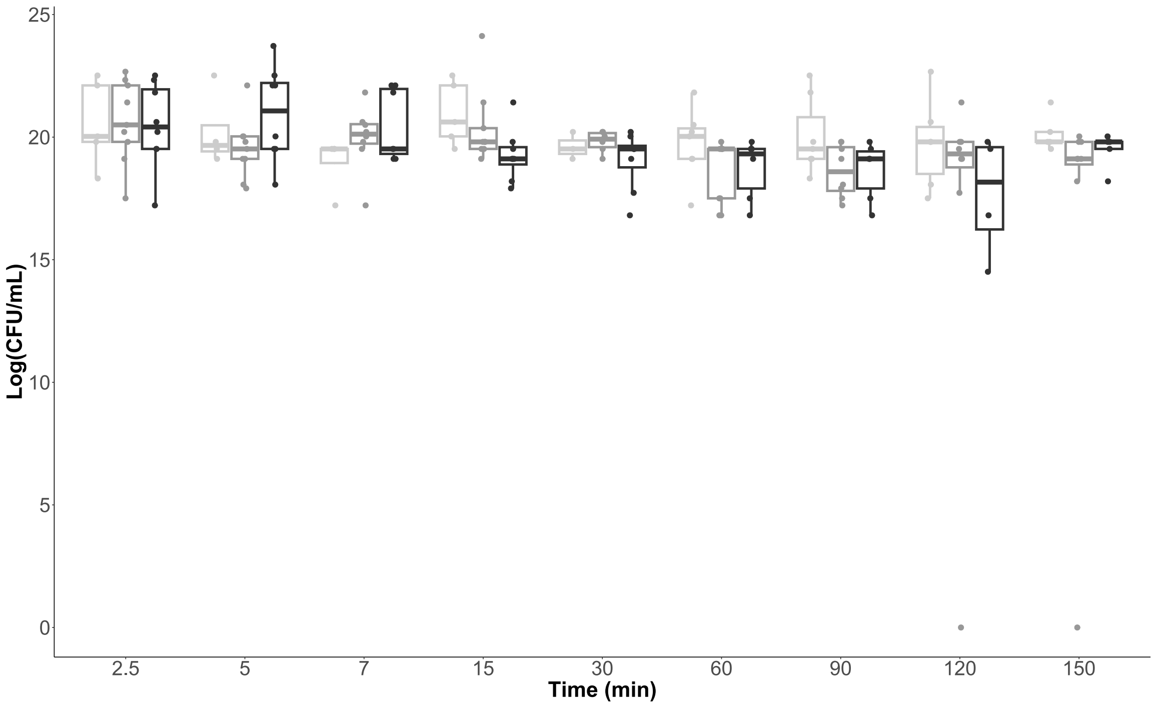


*Figure S2 Effect of pharamcokinatics without pexiganan challenge. Black boxes represent **Fast** dynamic, grey represents **Intermediate** dynamic, and light grey is **Slow** dynamic.*

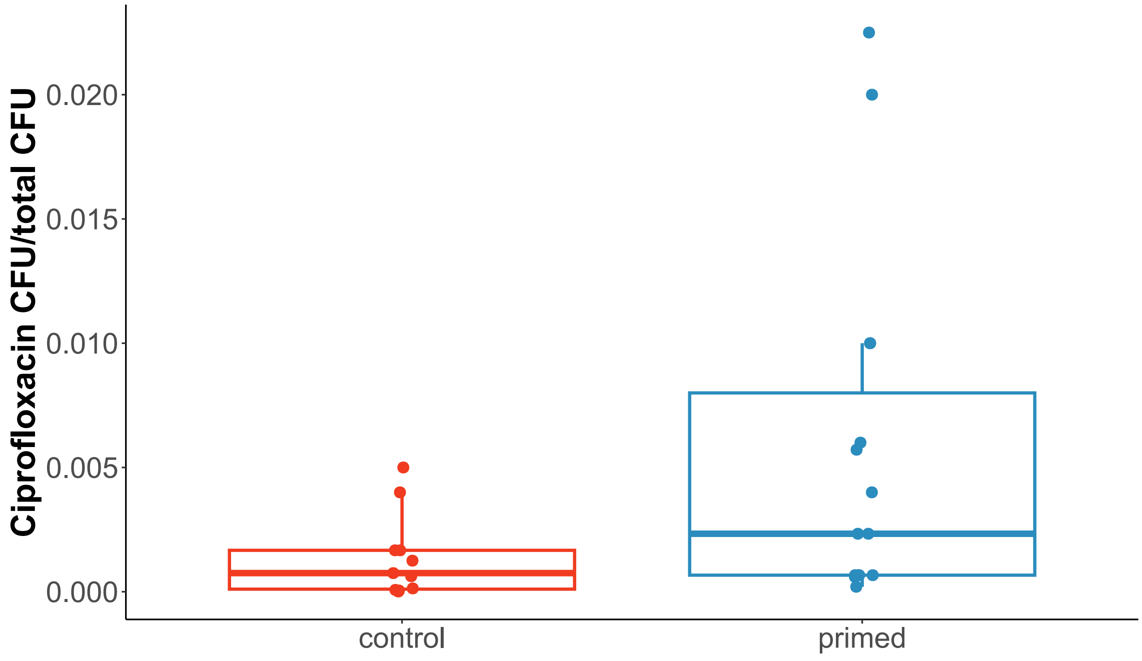


*Figure S3 Effect of priming on persister proportion, (Chisq = 4.3132, p = 0.03782).*

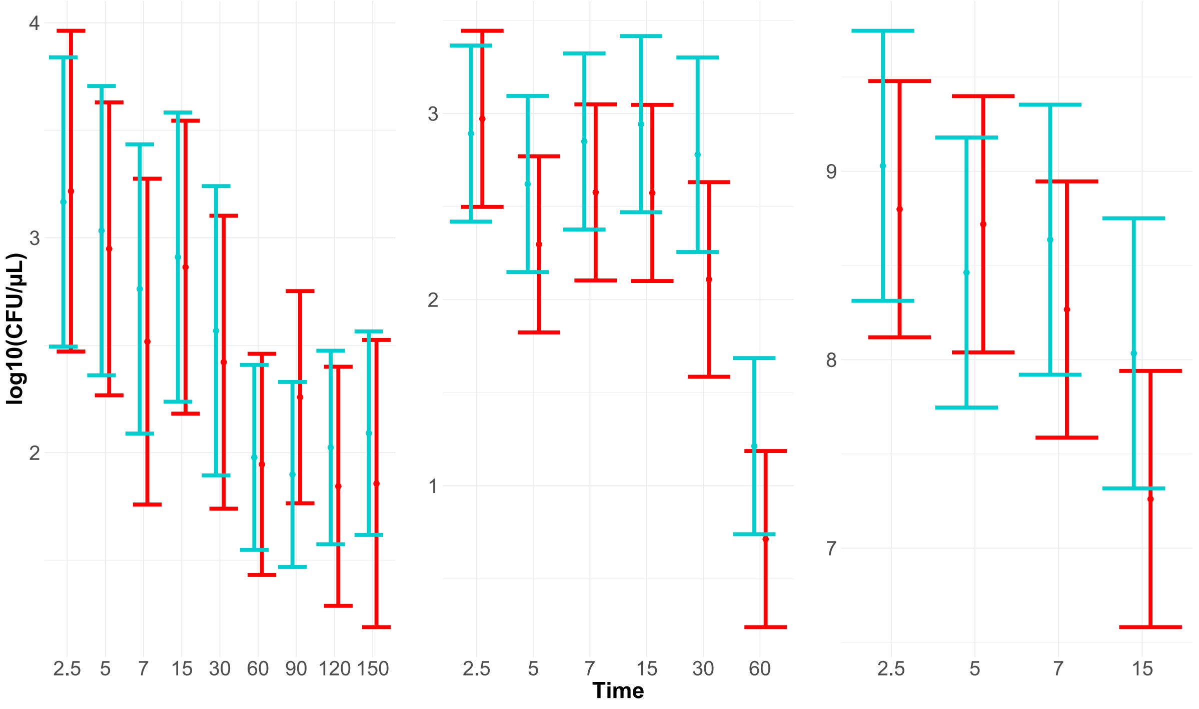


*Figure S4 Effect of time and treatment interaction on survival of primed/non-primed populations under dynamic application of pexiganan. From left to right, **Slow, Intermediate** and **Fast** application regimes are shown. The red bars represent the results of the non-primed bacterial cultures, whereas the blue bars represent the results of the primed cultures.*

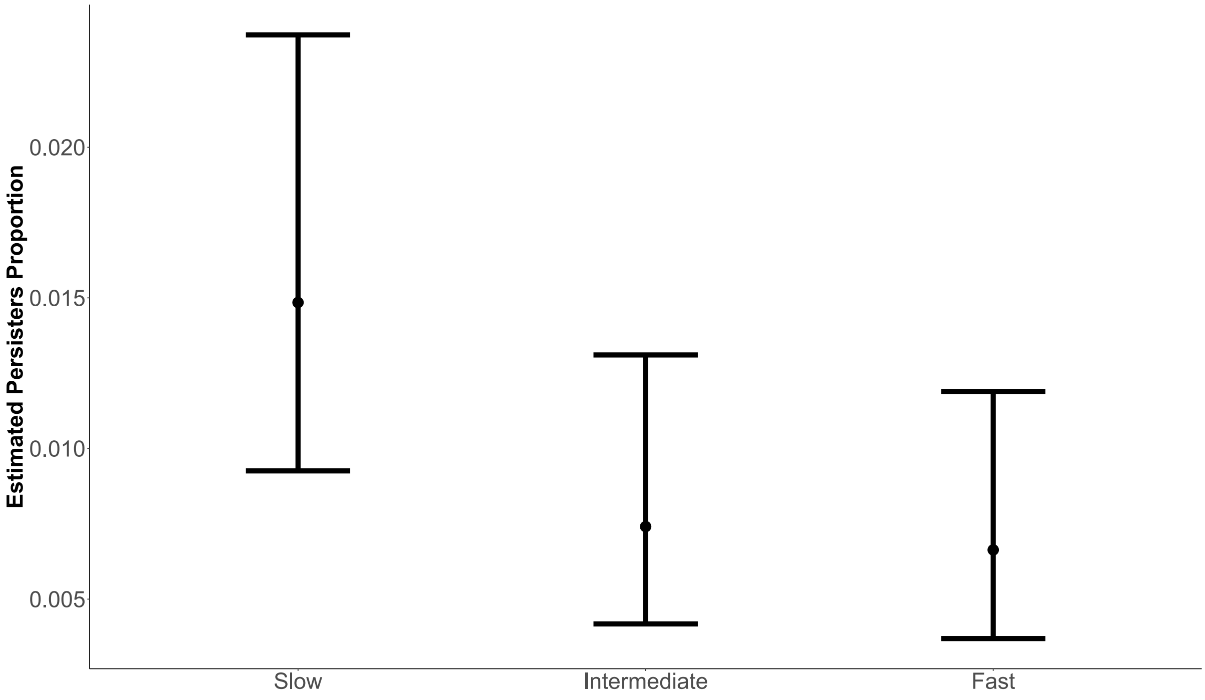


*Figure S5 Effect of application regime at 15 min on the proportion of persisters (Chisq = 7.7443, p <0.005).*
